# Negative regulation of uPAR activity by a GPI-specific phospholipase C

**DOI:** 10.1101/091272

**Authors:** Michiel van Veen, Elisa Matas-Rico, Koen van de Wetering, Daniela Leyton-Puig, Katarzyna M. Kedziora, Nicolai Sidenius, Kees Jalink, Anastassis Perrakis, Wouter H. Moolenaar

**Author notes:** Present address: Department of Dermatology and Cutaneous Biology, Sidney Kimmel Medical College at Thomas Jefferson University, Philadelphia, PA, USA. Present address: University of North Carolina, Chapel Hill Genetic Medicine Building, Chapel Hill, NC, USA. Equal first authorship (Michiel van Veen, Elisa Matas-Rico).

## Abstract

The urokinase receptor (uPAR) is a glycosylphosphatidylinositol (GPI)-anchored protein that promotes tissue remodeling, tumor cell adhesion, migration and invasion. uPAR mediates degradation of the extracellular matrix through protease recruitment and enhances cell adhesion, migration and signaling through vitronectin binding and interactions with integrins and other receptors. Full-length uPAR is released from the cell surface, but the mechanism and functional significance of uPAR release remain obscure. Here we show that transmembrane glycerophosphodiesterase GDE3 is a GPI-specific phospholipase C that cleaves and releases uPAR with consequent loss of the proteolytic and non-proteolytic activities of uPAR. In breast cancer cells, high GDE3 expression depletes endogenous uPAR resulting in a less transformed phenotype, correlating with higher survival probability in patients. Our results establish GDE3 as a negative regulator of the uPAR signaling network and, more generally, highlight GPI-anchor hydrolysis as a cell-intrinsic mechanism to alter cell behavior.

## Introduction

The urokinase-type plasminogen activator receptor (uPAR) is a central player in a complex signaling network implicated in a variety of remodeling processes, both physiological and pathological, ranging from embryo implantation to wound healing and tumor progression (Boonstra et al, 2011; Ferraris & Sidenius, 2013; Smith & Marshall, 2010). uPAR is a glycosylphosphatidylinositol (GPI)-anchored protein and hence lacks intrinsic signaling capacity. Instead, uPAR acts by binding two major ligands, namely the protease urokinase plasminogen activator (uPA) and the extracellular rmatrix (ECM) protein vitronectin (Ferraris & Sidenius, 2013; Madsen et al, 2007; Smith & Marshall, 2010). Through uPA binding, uPAR localizes plasmin generation to the cell surface and thereby promotes pericellular proteolysis and ECM degradation (Ferraris & Sidenius, 2013; Smith & Marshall, 2010). In addition, through vitronectin binding and functional interactions with integrins and growth factor receptors, uPAR activates intracellular signaling pathways leading to cytoskeletal reorganization, enhanced cell adhesion and motility and other features of tissue remodeling and cell transformation (Ferraris et al, 2014; Madsen et al, 2007; Smith & Marshall, 2010). As such, uPAR is a master regulator of extracellular proteolysis, cell motility and invasion. uPAR expression is elevated during inflammation and in many human cancers, where it often correlates with poor prognosis, supporting the view that tumor cells hijack the uPAR signaling system to enhance malignancy (Boonstra et al, 2011; Ferraris & Sidenius, 2013; Smith & Marshall, 2010).

It has long been known that full-length uPAR is released from the plasma membrane resulting in a soluble form (suPAR) (Pedersen et al, 1993; Ploug et al, 1992), which is detectable in body fluids and considered a marker of disease severity in cancer and other life-threatening disorders (Haupt et al, 2012; Hayek et al, 2016; Shariat et al, 2007; Sidenius et al, 2000; Stephens et al, 1999). Circulating suPAR is derived from activated immune and inflammatory cells (Ferraris & Sidenius, 2013; Smith & Marshall, 2010), and also from circulating tumor cells (Mustjoki et al, 2000). Locally produced suPAR might function as a ligand scavenger to confer negative feedback on uPAR (Smith & Marshall, 2010), but suPAR also can undergo proteolytic fragmentation possibly leading to new signaling activities (Montuori & Ragno, 2009). Yet, despite decades of research, the mechanism of uPAR release and its physiological implications have been elusive. A GPI-specific phospholipase D (GPI-PLD) (Scallon et al, 1991) has often been assumed to mediate the cleavage and release of GPI-anchored proteins, but this unique PLD does not function on native membranes and normally acts intracellularly during GPI biosynthesis (Mann et al, 2004).

A possible clue to the mechanism of uPAR release comes from recent studies showing that a member of the glycerophosphodiester phosphodiesterase (GDPD/GDE) family (Corda et al, 2014), termed GDE2, promotes neuronal differentiation by cleaving select GPI-anchored proteins, notably a Notch ligand regulator and heparan sulfate proteoglycans (Matas-Rico et al, 2016a; Matas-Rico et al, 2016b; Park et al, 2013). GDE2, along with GDE3 and GDE6, belongs to a GDE subfamily characterized by six-transmembrane-domain proteins with a conserved catalytic ectodomain (Figure 1a) (Corda et al, 2009; Matas-Rico et al, 2016a). GDE2’s closest relative, GDE3, accelerates osteoblast differentiation through an unidentified mechanism (Corda et al, 2009; Yanaka et al, 2003), while the function of GDE6 is unknown.

**Figure 1.**
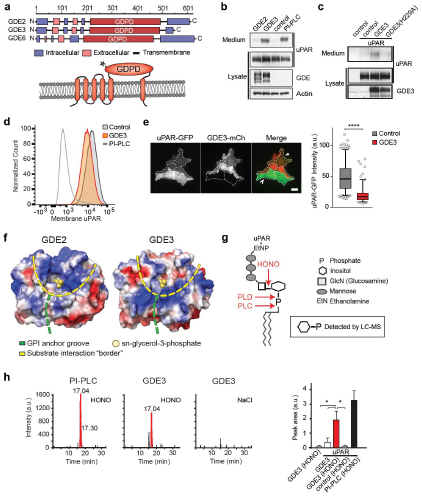
GDE3 is a GPI-PLC that cleaves and releases uPAR. **a**, GDE domain structure and transmembrane scheme of GDE3; asterisk, His229. **b**, Immunoblot analysis of uPAR release into the medium. HEK-uPAR cells transfected with empty vector (control), GDE2 or GDE3. PI-PLC served as positive control. **c**, GDE3(H229A) fails to release uPAR. **d, e**, uPAR disappears from the plasma membrane as measured by flow cytometry (**d**) and TIRF microscopy (**e**). Box plot shows uPAR-GFP intensity at the ventral membrane. **f**, Homology modeling of the GDE2 and GDE3 catalytic domains showing surface charge distributions (blue, positive; red, negative; green line, putative GPI-binding groove; yellow line, substrate-binding surface). The active site is indicated by glycerol-3-phosphate located at the template structure. **g**, GPI-anchor with cleavage sites indicated; HONO, nitrous acid. **h**, Representative LC-MS ion chromatograms (*m/z* 259.02-259.03); inositol 1-phosphate peaks in red. HONO-treated suPAR contains inositol 1-phosphate (n=3, mean ± SEM; \**P*<0.05 ANOVA).

Here we report that GDE3 is the long-sought phospholipase that cleaves and releases uPAR with consequent loss of its proteolytic and non-proteolytic activities.

## Results and Discussion

We set out to determine whether uPAR can be released by any of the three GDE family members, GDE2, GDE3 and GDE6. When expressed at relatively low levels in HEK293 cells, the GDEs localized to distinct microdomains at the plasma membrane, possibly representing clustered lipid rafts where GPI-anchored proteins normally reside (Maeda & Kinoshita, 2011), as well as to filopodia-like extensions (Figure 1 – figure supplement 1 and ref. (Matas-Rico et al, 2016a)). To assess GDE activity, we generated stable uPAR-expressing HEK293 cells (HEK-uPAR cells), expressed the respective GDEs and examined the appearance of suPAR in the medium, using bacterial phospholipase C (PI-PLC) as a control (Matas-Rico et al, 2016a). Strikingly, uPAR was readily released into the medium by GDE3, but not by GDE2 and GDE6 (Figure 1b and Figure 1-figure supplement 2a), thereby depleting the PI-PLC-sensitive uPAR pool at the plasma membrane (Figure 1d and Figure 1 – figure supplement 2b). A transmembrane version of uPAR (uPAR-TM) lacking the GPI moiety (Cunningham et al, 2003) was resistant to GDE3 attack, consistent with GDE3 acting through GPI-anchor hydrolysis (Figure 1 – figure supplement 2c). Furthermore, mutating putative active-site residue H229, corresponding to H233 in GDE2 (Matas-Rico et al, 2016a), abolished GDE3 activity (Figure 1c). Depletion of the GPI-anchored uPAR pool by GDE3 was most prominently observed at the ventral plasma membrane, as revealed by TIRF microscopy (Figure 1e).

To understand the structural basis of the marked GDE substrate selectivity, we constructed homology-based models of the globular α/β barrel GDPD domains, using I-TASSER (Yang et al, 2015). We reasoned that the catalytic domains must recognize not only the PI lipid moiety at the membrane-water interface, but also the attached protein. While GDE2 and GDE3 have a similar putative GPI-binding groove close to the active site, they show striking differences in their surface charge distribution, particularly at the putative substrate interaction surface (Figure 1f). It therefore seems likely that surface properties are a major determinant of selective substrate recognition by GDEs, a notion that should be validated by future structural studies.

We asked whether GDE3 cleaves uPAR in *cis* or in *trans*, or both. By mixing GDE3-expressing cells (lacking uPAR) with uPAR-expressing cells (lacking GDE3), we found that GDE3-expressing cells failed to release uPAR from the GDE3-deficient cell population (Figure 1 – figure supplement 2d). Thus, GDE3 cleaves uPAR on the same plasma membrane, not on adjacent cells. To determine whether GDE3 acts as a phospholipase C or D to hydrolyze the GPI-anchor, suPAR was immunoprecipitated from the medium of GDE3-expressing cells and treated with nitrous acid to cleave the glucosamine-inositol linkage in the GPI core (Figure 1g). Subsequent analysis by liquid chromatography-mass spectrometry (LC-MS) revealed that the acid-treated suPAR samples contained inositol 1-phosphate (Figure 1h). This result defines GDE3 as a GPI-specific PLC.

When compared to uPAR-deficient cells, HEK-uPAR cells showed markedly increased cell adhesion, loss of intercellular contacts and enhanced spreading with prominent lamellipodia formation on vitronectin, but not on fibronectin (Figure 2), features of a Rac-driven motile phenotype, in agreement with previous studies (Kjoller & Hall, 2001; Madsen et al, 2007). Cell spreading coincided with activation of focal adhesion kinase (FAK), indicative of integrin activation (Figure 2b). Strikingly, expression of GDE3 largely abolished the uPAR-induced phenotypes and cellular responses (Figure 2a-f). Catalytically dead GDE3(H229A) had no effect, neither had overexpressed GDE2 (Figure 2f and results not shown). Thus, by releasing uPAR from the plasma membrane, GDE3 suppresses the vitronectin-dependent activities of uPAR. Treatment of diverse cell types with suPAR-enriched conditioned media did not evoke detectable cellular responses, supporting the notion that suPAR is biologically inactive, at least under cell culture conditions.

**Figure 2.**
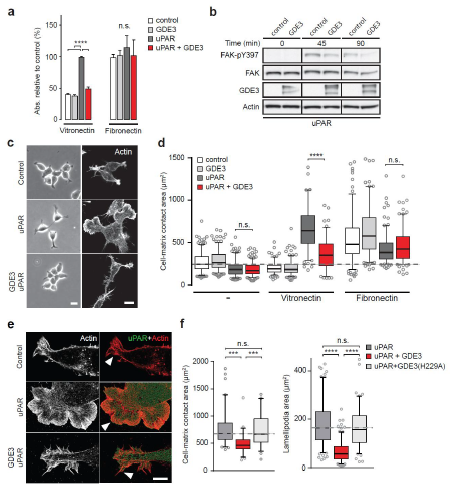
GDE3 inhibits uPAR activity in HEK-uPAR cells. **a**, uPAR confers increased adhesion to vitronectin, but not fibronectin, which is prevented by GDE3 expression (n=3, mean ± SEM), *\*\*\*\*P* <0.0001; n.s., not significant. **b**, GDE3 inhibits FAK activation during cell adhesion. FAK activity was assayed at the indicated times after plating. **c**, GDE3 inhibits uPAR-induced cell spreading, scattering and lamellipodia formation on vitronectin (bar, 10 μm). **d**, Cell-matrix contact area of cells expressing the indicated constructs plated on either fibronectin or vitronectin. Box plots show the mean of three independent experiments. Dotted line represents the mean of control cells. **e,f**, uPAR-induced lamellipodia on vitronectin disappear upon GDE3 expression, as shown by super-resolution microscopy (**e**) (bar, 5 μm) and quantified in (**f**). GDE3(H229A) fails to affect lamellipodia formation. \*\*\**P*<0.0005; \*\*\*\**P*<0.00005 of ANOVA.

We next assessed the impact of GDE3 on endogenous uPAR activity in MDA-MB-231 in triple-negative breast cancer cells. These cells express relatively high levels of uPAR (Figure 3a and Figure 3 - figure supplement 1a) and its ligand uPA (LeBeau et al, 2013), thus forming an autocrine signaling loop. GDE3 expression is relatively low in human cancer cell lines, including breast cancer lines (n=51) (Barretina et al, 2012). We overexpressed GDE3 in MDA-MB-231 cells, confirmed its localization to the plasma membrane (Figure 3 - figure supplement 1b) and examined its impact. Wild-type cells adopted a motile phenotype on vitronectin (but not on fibronectin), as evidenced by increased cell spreading with marked lamellipodia formation (Figure 3b-d), strongly reminiscent of a uPAR-regulated phenotype. Overexpression of GDE3 led to a marked loss of uPAR protein from the cells, without affecting uPAR mRNA levels (Figure 3a), and abolished the vitronectin-dependent phenotype (Figure 3b-d). No effect was observed upon GDE2 overexpression (data not shown). To confirm that GDE3 acts through uPAR depletion, we expressed non-cleavable uPAR-TM to compete out endogenous uPAR, and found that GDE3-induced cell spreading on vitronectin was largely inhibited (Figure 3c). Furthermore, RNAi-mediated knockdown of uPAR (Figure 3e) gave rise to the same phenotype as GDE3 overexpression, namely reduced cell adhesion, spreading and lamellipodia formation on vitronectin (Figure 3f-h). In long-term assays, wild-type MDA-MB-231 cells showed marked scattering, indicative of increased cell motility with loss of intercellular contacts (Figure 4a). Again, GDE3 overexpression mimicked uPAR depletion in greatly reducing both cell motility and clonogenic potential (Figure 4a,b).

**Figure 3.**
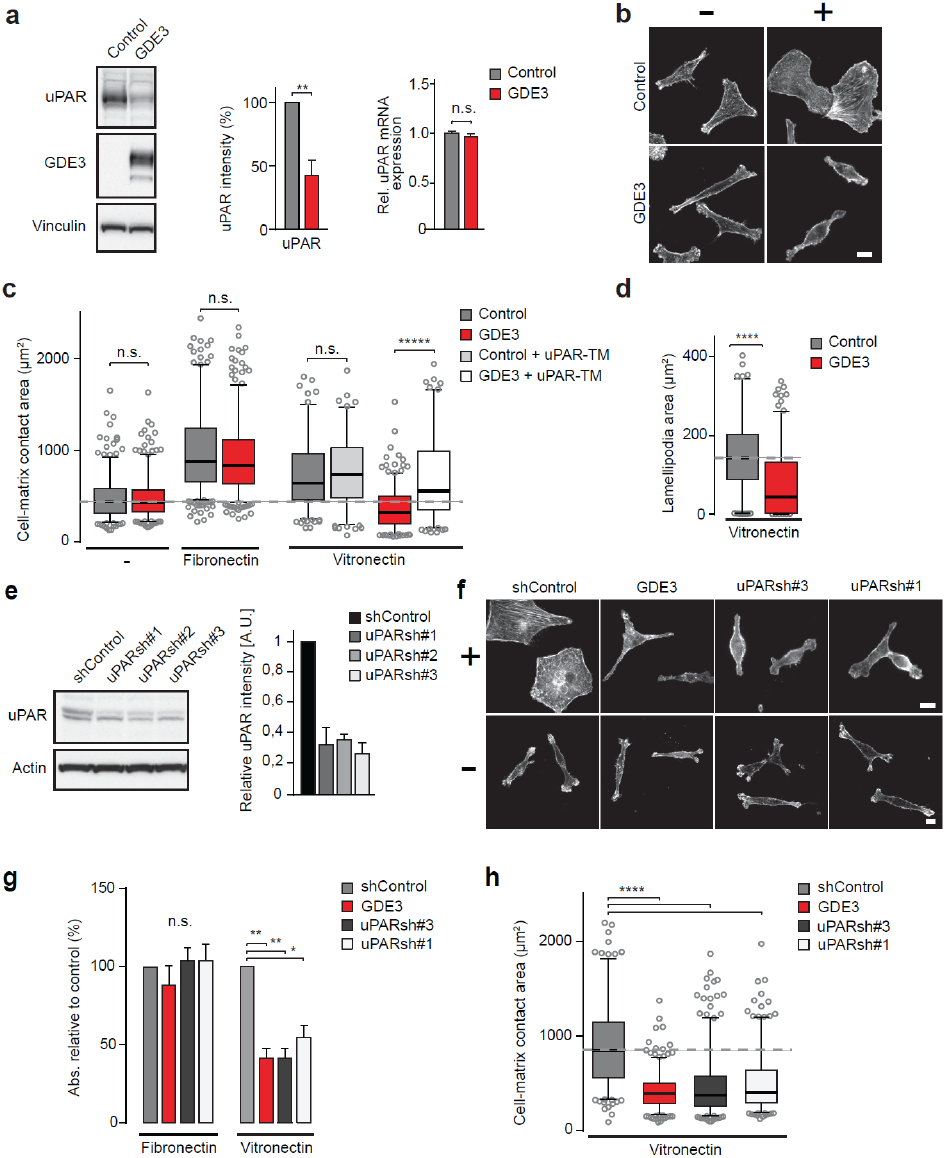
GDE3 depletes uPAR and suppresses the vitronectin-dependent phenotype of MDA-MB-231 breast cancer cells. **a**, GDE3 overexpression decreases uPAR protein but not mRNA expression, as shown by immunoblot and qPCR, respectively (n=3, mean ± SEM). **b**, Confocal images showing that GDE3 inhibits cell spreading and lamellipodia formation on vitronectin (+). Bar, 10 μm. **c,d**, GDE3 reduces cell spreading (**c**) and lamellipodia formation (**d**) on vitronectin, whilst non-cleavable uPAR-TM inhibits GDE3 activity (**c**). **e**, Immunoblot analysis of uPAR knockdown; maximum knockdown was achieved by hairpins #1 and #3. **f,g,h**, GDE3 overexpression phenocopies uPAR knockdown in cells on vitronectin (+); bar, 10 μ (**f**).

**Figure 4.**
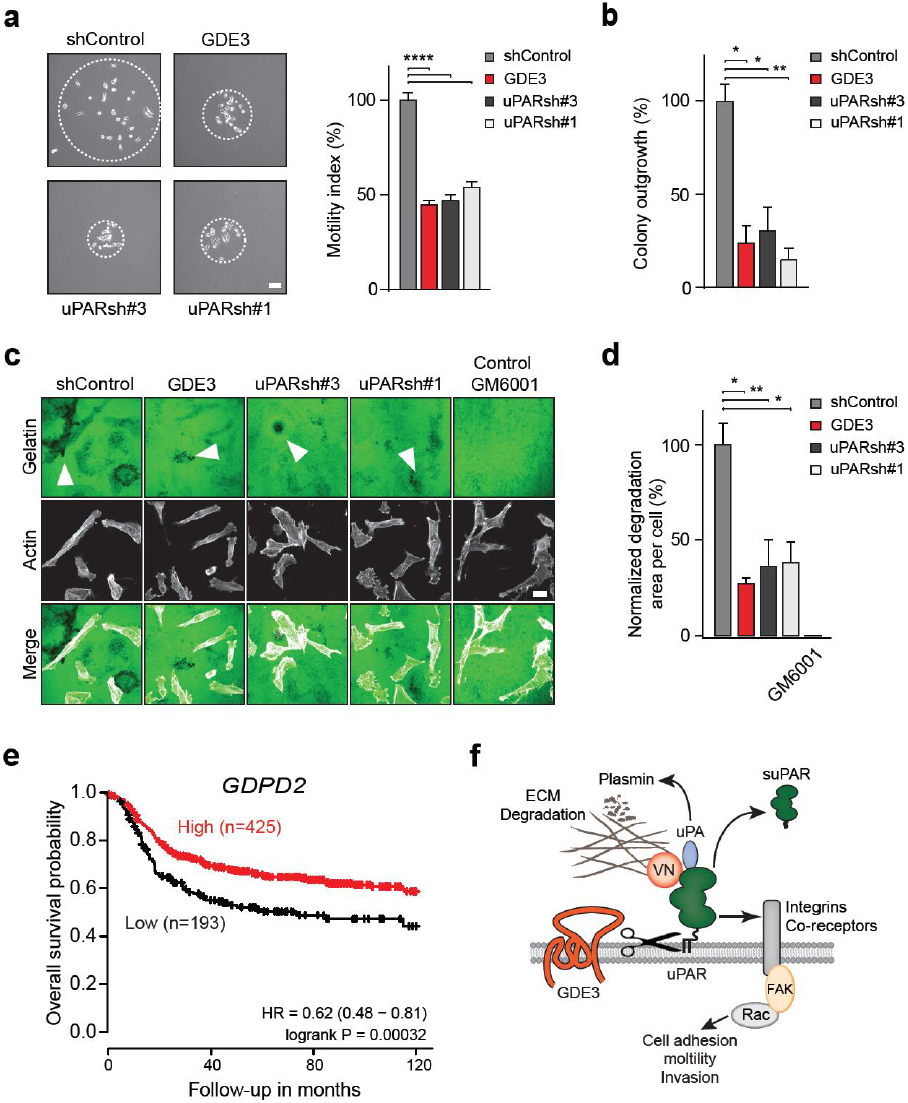
GDE3 expression attenuates the uPAR-dependent transformed phenotype of breast cancer cells, correlating with higher survival probability. **a,b**, GDE3 overexpression in MDA-MB-231 cells mimics uPAR knockdown in inhibiting cell scattering (**a**) and colony formation (**b**). In (**a**), MDA-MB-231 cells grow out as scattered colonies; bar, 100 μm. (n=3, mean ±SEM ** * *P*<0.0005). In (**b**), colonies were counted after 14 days. * *P*<0.05 of ANOVA. **c,d**, Degradation of a gelatin matrix (bar, 10 μm). Arrows point to black spots where gelatin was degraded. Metalloprotease inhibitor GM6001 was used as a control (**c**). Degradation was quantified (**d**) at 20 hrs after plating (mean ± SEM **** *P*<0.0005 of ANOVA). (**e**) High *GDPD2* expression significantly correlates with higher survival rate in triple-negative breast cancer patients (n=618). HR, hazard ratio with 95% confidence interval. Analysis based on microarray data (www.kmplot.com). (**f**) GDE3 cleaves and releases uPAR leading to loss of uPAR function.

Having shown that GDE3 suppresses the non-proteolytic activities of uPAR, we next examined how GDE3 affects uPAR-driven proteolytic matrix degradation. Also in this case, GDE3 overexpression mimicked uPAR knockdown in inhibiting the degradation of a gelatin matrix (Figure 4c,d). We therefore conclude that GDE3 attenuates the transformed phenotype of uPAR-positive breast cancer cells through loss of functional uPAR. Interestingly, high expression of GDE2 (encoded by *GDPD2*) was found to be associated with improved relapse-free survival in breast cancer, particularly in triple-negative subtype patients (Figure 4e). No such correlation was found for GDE2. These results are consistent with the cell-based data; however, we cannot rule out that GDE3 may cleave additional GPI-anchored substrates whose functional loss could contribute to positive clinical outcome.

In conclusion, our results reveal GDE3 as the long-sought cell-intrinsic GPI-PLC that releases uPAR, and establish GDE3 as a negative regulator of the uPAR signaling network (Figure 4f). Thus, depending on its expression level, GDE3 is predicted to downregulate normal uPAR-dependent remodeling processes. Indeed, upregulated GDE3 accelerates osteoblast differentiation (Corda et al, 2009; Yanaka et al, 2003) in a manner resembling the uPAR knockout phenotype (Furlan et al, 2007). Furthermore, a striking >200-fold upregulation of *GDPD2* is observed during blastocyst formation (Munch et al, 2016), implicating GDE3 in the invasion of pre-implantation embryos, a known uPA/uPAR-dependent process (Multhaupt et al, 1994). Although correlative, these results support the view that GDE3 is upregulated to downregulate uPAR activity. The present findings also imply that circulating suPAR should be regarded as a marker of GDE3 activity, not necessarily reflecting uPAR expression levels.

It will now be important to determine how GDE3 expression and activity are regulated and, furthermore, to explore the substrate selectivity of GDEs in further detail. When regarded in a broader context, the present and previous findings (Matas-Rico et al, 2016a; Matas-Rico et al, 2016b; Park et al, 2013) support the view that vertebrate GDEs have evolved to modulate key signaling pathways and alter cell behavior through selective GPI-anchor cleavage.

## Materials and Methods

### Cell culture and materials

HEK293, MDA-MB-231 and MCF-7 cells, obtained from the ATCC, were grown in Dulbecco’s modified Eagle’s medium (DMEM) supplemented with 10% fetal bovine serum (FBS) and antibiotics at 37 °C under 5% CO__2__. Antibodies used: anti-mCh and anti-GFP, home-made; anti-Flag, M2, anti-Vinculin and β-Actin (AC-15) from Sigma; anti-uPAR (MAB807) from R&D systems; anti-FAK(pTyr397) from Thermo Fisher. Vitronectin, fibronectin, inositol 1-phosphate (dipotassium salt) and inositol were purchased from Sigma-Aldrich. *B. cereus* PI-PLC was from Molecular Probes. Phalloidin red (actin-stain^™^ 647 phalloidin) and green (actin-stain^™^ 488 phalloidin) were from Cytoskeleton. GM6001 was from Millipore.

### Expression vectors

GDE3 cDNA was amplified by PCR and subcloned into a pcDNA3-HA plasmid using AflII/HpaI (PCR product) and AflII/EcoRV (plasmid) restriction sites. GDE3 was recloned into a pcDNA3(-mCherry) construct by PCR amplification with restriction sites PmI1/Xba1, followed by vector digestion using EcoRV/Xba1. All constructs were tagged on the C-terminus unless otherwise stated. Mutant GDE3(H229A) was generated by amplification with oligos containing the mutation, followed by Dpn1 digestion of the template. The viral plasmids (pBABE-GDE3-mCh, pBABE-uPAR-GFP) were constructed by subcloning the GDE3-pcDNA3 and uPAR-GFP-pEGFP-N1 into a pBABE plasmid. GDE3-mCherry-pcDNA3 was cut using PmeI followed by digestion of the pBABE backbone with SnaBI. uPAR-GFP-pEGFP-N1 was cut with BglII and HpaI and ligated into the pBABE vector, digested with BAMHI and SnaB1. Constructs uPAR-GFP, uPAR-FLAG and uPAR-TM were previously described(Cunningham et al, 2003). Human GDE2 cDNA was subcloned as described(Matas-Rico et al, 2016a). GDE2-FLAG constructs were provided by Dr. Sungjin Park.

### Transfections and knockdown

Cells stably expressing uPAR-GFP or GDE3-mCherry were generated using retroviral transduction and subsequent selection with puromycin. Transient transfections were done using the calcium phosphate protocol or XtremeGene 9 agent (Roche). Stable uPAR knockdown in MDA-MB-231 cells was achieved using shRNAs in a lentiviral pLKO vector; five shRNAs from three RC human shRNA library were tested: TRCN0000052637, TRCN0000052636, TRCN0000052634, TRCN0000052633 and TRCN0000052635. The latter two were used for experiments; sequences: CCGGCCCATGAATCAATGTC TGGTACTCGAGTACCAGACATTGATTCATGGGTTTTTG and CGGGCTTGAAGA TCACCAGCCTTACTCGAGTAAGGCTGGTGATCTTCAAGCTTTTTG, respectively. For virus production, HEK293T cells were transiently transfected using calcium phosphate, and virus particles were collected 48 hr thereafter. uPAR knockdown cells were selected in medium containing 2μg/ml puromycin.

### Liquid chromatography-mass spectrometry (LC-MS)

To determine the inositol phosphate content of cleaved uPAR, suPAR was immuno-precipitated from HEK293 cell conditioned medium using anti-GFP beads (ChromoTek). To remove inositol phosphate from suPAR, the beads were treated with 0.1M acetate buffer (pH 3.5) and subsequently with 0.5M NaNO__2__ or 0.5M NaCl (Control) for 3 hrs as previously described (Mehlert & Ferguson, 2009). Inositol phosphate-containing samples were preprocessed by adding methanol to a final concentration of 70% and shaken at 1000 RPM at room temperature for 10 minutes. Following centrifugation (20,400 × *g* at 4 °C for 10 min), the supernatant was evaporated to dryness in a Speedvac at room temperature. The dried extracts were reconstituted in 50 mM ammonium acetate (pH 8.0), centrifuged (20,400 × *g* at 4 °C for 10 min) and transferred to autosampler vials. Liquid chromatography (LC) was performed using a Dionex Ultimate 3000 RSLCnano system (Thermo Fisher Scientific). A volume of 5 μl was injected on a Zorbax HILIC PLUS column (150 × 0.5 mm, 3.0 μm particles) maintained at 30 °C. Elution was performed using a gradient: (0-5 min, 20% B; 5-45 min 20-100% B; 45-50 min 100% B; 50-50.1 min 20% B; 50.1-60 min 20% B) of 100% acetonitrile (mobile phase A) and 50 mM ammonium acetate adjusted to pH 8.0 with ammonium hydroxide (mobile phase B) at a flow rate of 15 μl/min. Inositol 1-phosphate was detected with an LTQ-Orbitrap Discovery mass spectrometer (Thermo Fisher Scientific) operated in negative ionization mode scanning from *m/z* 258 to 260 with a resolution of 30,000 FWHM. Electrospray ionization was performed with a capillary temperature set at 300 °C and the sheath, auxiliary and sweep gas flow set at 17, 13 and 1 arbitrary units (AU), respectively. Setting for Ion guiding optics were: Source voltage: 2.4 kV, capillary voltage: −18 V, Tube Lens: −83 V, Skimmer Offset: 0 V, Multipole 00 Offset: 5 V, Lens 0: 5 V, Multipole 0 Offset: 5.5 V, Lens 1: 11 V, Gate Lens Offset: 68 V, Multipole 1 Offset: 11.5 V, Front Lens: 5.5 V. Data acquisition was performed using Xcalibur software (Thermo Fisher Scientific). Reference inositol phosphate (Sigma) was used to determine the retention on the Zorbax HILIC plus column. After applying Xcalibur’s build-in smoothing algorithm (Boxcar, 7), extracted ion chromatograms *(m/z* 259.02-259.03) were used to semi-quantitatively determine inositol phosphate levels.

### Cell adhesion and spreading

48-well plates were coated overnight at 4°C with fibronectin (10 μg/ml) or vitronectin (5 μg/ml), or left untreated. Thereafter, plates were blocked for 2 hr at 37°C using 0.5% BSA in PBS. Cells were washed and harvested in serum-free DMEM supplemented with 0.1% BSA. Equal numbers of cells were seeded and allowed to adhere for 1 hr. Non-adherent cells were washed away using PBS. Attached cells were fixed with 4% paraformaldehyde (PFA) for 10 min, followed by washing and staining with Crystal violet (5 mg/ml in 2% ethanol) for 10 min. After extensive washing, cells were dried and lysed in 2% SDS for 30 min. Quantification was done by measuring absorbance at 570 nm using a plate reader. For cell-matrix contact area and lamellipodia measurements, coverslips were coated overnight with fibronectin or vitronectin and washed twice with PBS. Cells were trypsinized, washed and resuspended in DMEM and left to adhere and spread for 4 hr. After fixation (4% PFA) and F-actin staining with phalloidin, images were taken using confocal microscopy. Cell and lamellipodia area was quantified using an ImageJ macro.

### Cell scattering and colony formation

Cell scattering was determined as described (LeBeau et al, 2013). In brief, single MDA-MB-231 cells were allowed to grow out as colonies, and the area covered by the scattered colonies (colony size) was measured at 6 days after plating. For measuring colony outgrowth, 500 cells were plated and colonies were counted after 14 days.

### Matrix degradation

Coverslips were coated with OG-labelled gelatin (Invitrogen) supplemented with vitronectin (5 μg/ml). About 100.000 cells per coverslip were seeded in DMEM with 10% FCS. After 20 hrs, cells were washed, fixed with 4% PFM and stained with phalloidin-Alexa647 (Invitrogen). Gelatin degradation was determined from confocal images of >15 randomly chosen fields of view per coverslip (testing at least two coverslips / condition on two separate days: total four coverslips per condition). The images were randomized and the area of degradation was normalized to the total area of cells or to the number of cells.

### RT-qPCR

Total RNA was isolated using the GeneJET purification kit (Fermentas). cDNA was synthesized by reverse transcription from 2 μg RNA with oligodT 15 primers and SSII RT enzyme (Invitrogen). Relative qPCR was measured on a 7500 Fast System (Applied Biosystems) as follows: 95 °C for 2 min followed by 40 cycles at 95 °C for 15 sec followed by 60 °C for 1 min. 200 nM forward and reverse primers, 16 μl SYBR Green Supermix (Applied Biosystems) and diluted cDNA were used in the final reaction mixture. GAPDH was used as reference gene and milliQ was used as negative control. Normalized expression was calculated following the equation NE=2^(Ct target-Ct reference)^. Primers used: GDE3, forward TCAGCAGGACCACGAATGTA, reverse GCTGCAGCTTCCTCCAATAG; uPAR, forward AATGGCCGCCAGTGTTACAG, reverse CAGGAGACATCAATGTGGTTC; Cyclophilin, forward CATCTGCACTGCCAAGACTGA, reverse TTGCCAAACACCACATGCTT.

### Western blotting

For Western blotting, cells were washed with cold PBS, lysed in RIPA buffer supplemented with protease inhibitors and spun down. Protein concentration was measured using a BCA protein assay kit (Pierce) and LDS sample buffer (NuPAGE, Invitrogen) was added to the lysate or directly to the medium. Equal amounts were loaded on SDS-PAGE pre-cast gradient gels (4-12% Nu-Page Bis-Tris, Invitrogen) followed by transfer to nitrocellulose membrane. Non-specific protein binding was blocked by 5% skimmed milk in TBST; primary antibodies were incubated overnight at 4° C in TBST with 2% skimmed milk. Secondary antibodies conjugated to horseradish peroxidase (DAKO, Glostrup, Denmark) were incubated for 1 hr at room temperature; proteins were detected using ECL Western blot reagent.

### Microscopy

Cells cultured on 24 mm, #1,5 coverslips were washed and fixed with 4% PFA, permeabilized with 0.1% Triton X-100 and blocked with 2% BSA for 1 hr. Incubation with primary antibodies was done for 1 hr followed by incubation with Alexa-conjugated antibodies or Phalloidin for 45 min at room temperature. For confocal microscopy, cells were washed with PBS, mounted with Immnuno-MountTM (Thermo Scientific) and visualized on a LEICA TCS-SP5 confocal microscopy (63 × objective). Super-resolution imaging was done using a SR-GSD Leica microscope equipped with an oxygen scavenging system, as previously described (Matas-Rico et al, 2016a). In short, 15000 frames were taken in TIRF mode, at 10 ms exposure time. After post image analysis, movies were analyzed and corrected using the ImageJ plugin Thunderstorm (http://imagej.nih.gov/ij/) followed by correction with an ImageJ macro using the plugin Image Stabilizer. For Total Internal Reflection (TIRF) microscopy, HEK293 cells stably expressing UPAR-GFP and transiently transfected with GDE3-mCherry were imaged using a Leica AM TIRF MC microscope with a HCX PL APO 63x, 1.47 NA oil immersion lens. Excitation was at 488 and 561 nm and detection of fluorescence emission was by a GR filter cube (Leica). Before each experiment, automatic laser alignment was carried out and TIRF penetration depth was set to 200 nm. Data were acquired at 500 ms frame rate.

### Flow cytometry

HEK293 cells stably expressing uPAR-GFP were left untreated, treated with PI-PLC or transiently transfected with GDE3-mCherry. Cells were trypsinized, blocked in 2%BSA and stained with rabbit anti-GFP primary antibody followed by AlexaFluor-647 coupled anti-rabbit secondary antibody. Cells were analyzed using a BD LSR Fortessa flow cytometer.

### Statistical analysis

For all single comparisons, a two tailed unpaired Student’s t-test was used; for multiple comparisons an ordinary ANOVA with Tukey’s test. A *P* value <0.05 was considered statistically significant. Error bars shown in the bar diagrams were calculated as the standard error of the mean (SEM); whiskers in the box plots depict 95% confidence intervals.

## Acknowledgements

We thank Sungjin Park and Shantini Sockanathan (Johns Hopkins University School of Medicine, Baltimore, USA) for reagents and fruitful discussions. This work was supported by the Netherlands Organisation for Scientific Research (NWO) and the Dutch Cancer Society (KWF).

---

**Figure 1 – figure supplement 1.**
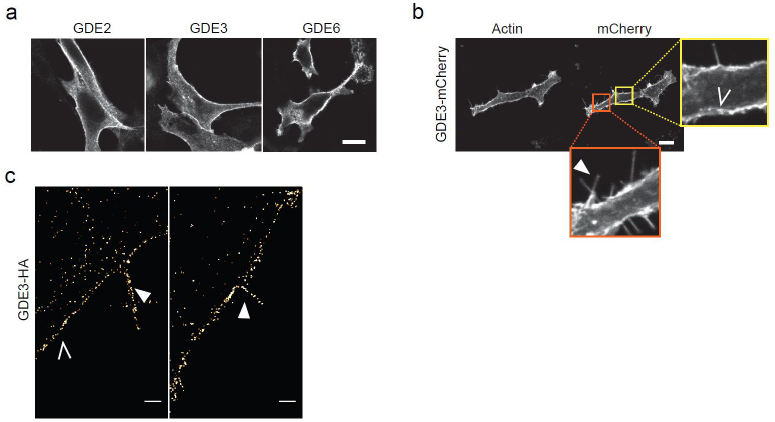
GDE subcellular localization. **a**, Confocal images of HEK293 cells expressing GDE2-HA, GDE3-HA or GDE6-FLAG; bar, 10 μm. **b,c**, GDE3 localizes to distinct microdomains (yellow square, open arrows) and filopodia-like extensions (orange square, solid arrow), as visualized by confocal (b) and super-resolution (c) microscopy. Phalloidin was used to visualize F-actin. Bars, 10 μm (a, b) and 1 μm (c), respectively.

**Figure 1 – figure supplement 2.**
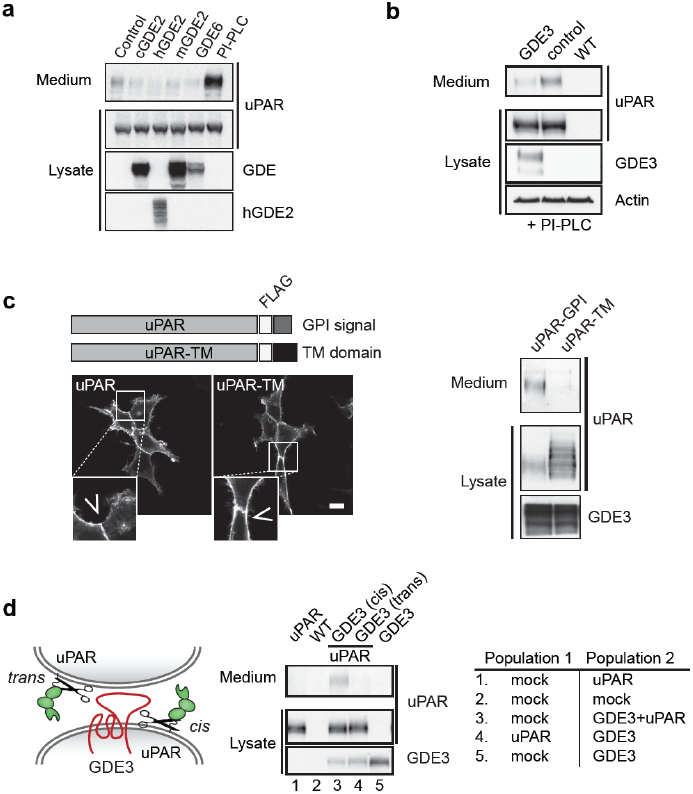
Induction of uPAR release from HEK293-uPARcells. **a**, Immunoblot analysis showing that vertebrate GDE2 (chicken, mouse and human) and GDE6 fail to release uPAR. PI-PLC was used as positive control. **b**, Immunoblot analysis showing that GDE3 depletes the membrane-anchored uPAR pool. HEK-uPAR cells expressing GDE3 or empty vector (control) were treated with PI-PLC. Right lane refers to wild-type (WT) HEK293 cells. **c**, uPAR-TM, containing the transmembrane domain of EGFR, is properly expressed at the plasma membrane (bar, 10 μm), but is not released by GDE3 as shown by immunoblotting using anti-FLAG antibody. **d**, (Left) Scheme showing uPAR cleavage in *cis* or *trans*. (Right) GDE3-expressing HEK-uPAR cells were mixed with a GDE3-deficient cell population, as indicated. Immunoblot analysis of uPAR in medium and cell lysates indicates that GDE3 acts in *cis;* mock refers to empty vector-transfected cells.

**Figure 3 – figure supplement 1.**
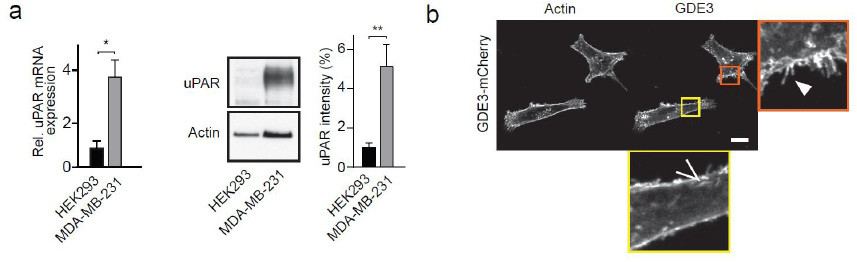
uPAR expression and GDE3 localization in MDA-MB-231 cells. **a**, Expression of uPAR protein and mRNA in MDA-MB-231 versus HEK293 cells. * *P*<0.05; ** *P* < 0.001 of t-test. **b**, Membrane localization of GDE3-mCh to distinct microdomains and filopodia-like extensions; bar, 10 μm.

